# Resource selection by the Endangered Arabian tahr: identifying critical habitats for conservation and climate change adaptation

**DOI:** 10.1101/2021.01.26.428299

**Authors:** S Ross, H. Rawahi, M.H. Jahdhami

## Abstract

The Arabian tahr is an Endangered mountain ungulate endemic to the Hajar Mountains of Arabia. The Arabian tahr population is in decline and threats to tahr habitat are intensifying, in addition new potential challenges from climate change are emerging. Fundamental to future conservation planning is understanding tahr habitat selection patterns, so we can prioritise habitat protection, and understand how habitat may be used to provide thermal refuge and allow adaptation to climate change impacts. We used GPS collars and resource selection functions to characterise Arabian tahr habitat preferences in Wadi Sareen Nature Reserve, Oman. We found tahr habitat selection was dependent on scale, sex and season. Vegetation resources were only selected at the smallest scales of selection and avoided at other scales. Habitat providing low heat load and thermal refuge were intensely selected at small and medium scales, by both sexes and in both seasons, suggesting the importance of thermal refuges in facilitating thermoregulation. Higher elevations, steep slopes and rugged habitats were selected across all scales tested here, and in previous landscape-scale studies, indicating the fundamental importance of these habitats in supporting Arabian tahr populations. Our results identified critical habitats required to sustain Arabian tahr, and demonstrated the importance of thermal refuges to species living in the hot climates such as the Arabian Peninsula. Given the accessibility of habitat layers, and ease in which the identified habitats can be mapped using a geographical information system, understanding the habitat selection of tahr and other species is a crucial step to increasing conservation management capacity of threatened species. Given our uncertainty of how to conserve wildlife under future climate change, understanding the availability and distribution of wildlife habitat is an important baseline from where we can plan, connect and preserve the resources necessary for wildlife conservation.

## Introduction

Since the 1970s following the discovery of large oil reserves, the Arabian Peninsula has seen an astonishing rate of economic and industrial development (Zahlan, 2016). Despite the many benefits of economic development, there have also been consequences and impacts to the region’s unique assemblage of arid land biodiversity. Although development in Oman has progressed more slowly than the rest of the region, there has still been large-scale loss and fragmentation of natural habitats (Ross et al., 2020) which has transformed the countries natural landscapes. Wildlife poaching and exploitation is also an ongoing issue in Oman (Giangaspero and Ghafri, 2014; Hikmani et al., 2015) which has had large impacts on populations of native fauna. To help limit the loss of biodiversity there has been a push by the Omani government to address conservation issues and build knowledge and conservation capacity, particularly for Endangered species such as the Arabian leopard (*Panthera pardus nimr*) and the Arabian tahr (*Arabitragus jayakari*; Burton et al., 2013).

The Arabian tahr is an elusive mountain goat endemic to northern Oman and western United Arab Emirates. The species is rare and seldom seen, frequenting the steep, inaccessible cliffs of the 700 km Hajar Mountain range (Fig. 1). Although the tahr is listed as Endangered in the IUCN Red List and is among the most vulnerable species in the Arabian Peninsula, it remains one of the world’s least studied ungulates. Our poor understanding restricts our ability to protect and manage the remaining tahr populations despite increasing disturbance within its Hajar Mountain range from infrastructural developments and resource extraction. Large-scale research has highlighted that tahr exist at a very low density across most of its range and only occupy an estimated 23.9% of their former distribution (Ross et al., 2017). They also face threats from poaching, habitat loss and fragmentation, population fragmentation, and displacement and disease risk from domestic livestock (Ross et al., 2019; 2020). Climate change is also expected to impact Arabian tahr, extreme temperatures in the region already reach up to 50°C in the summer, and increasing average and extreme temperatures are forecast due to climate change in the Arabian Peninsula (Pal and Eltahir, 2016; Varela et al., 2020).

**Figure 1:**
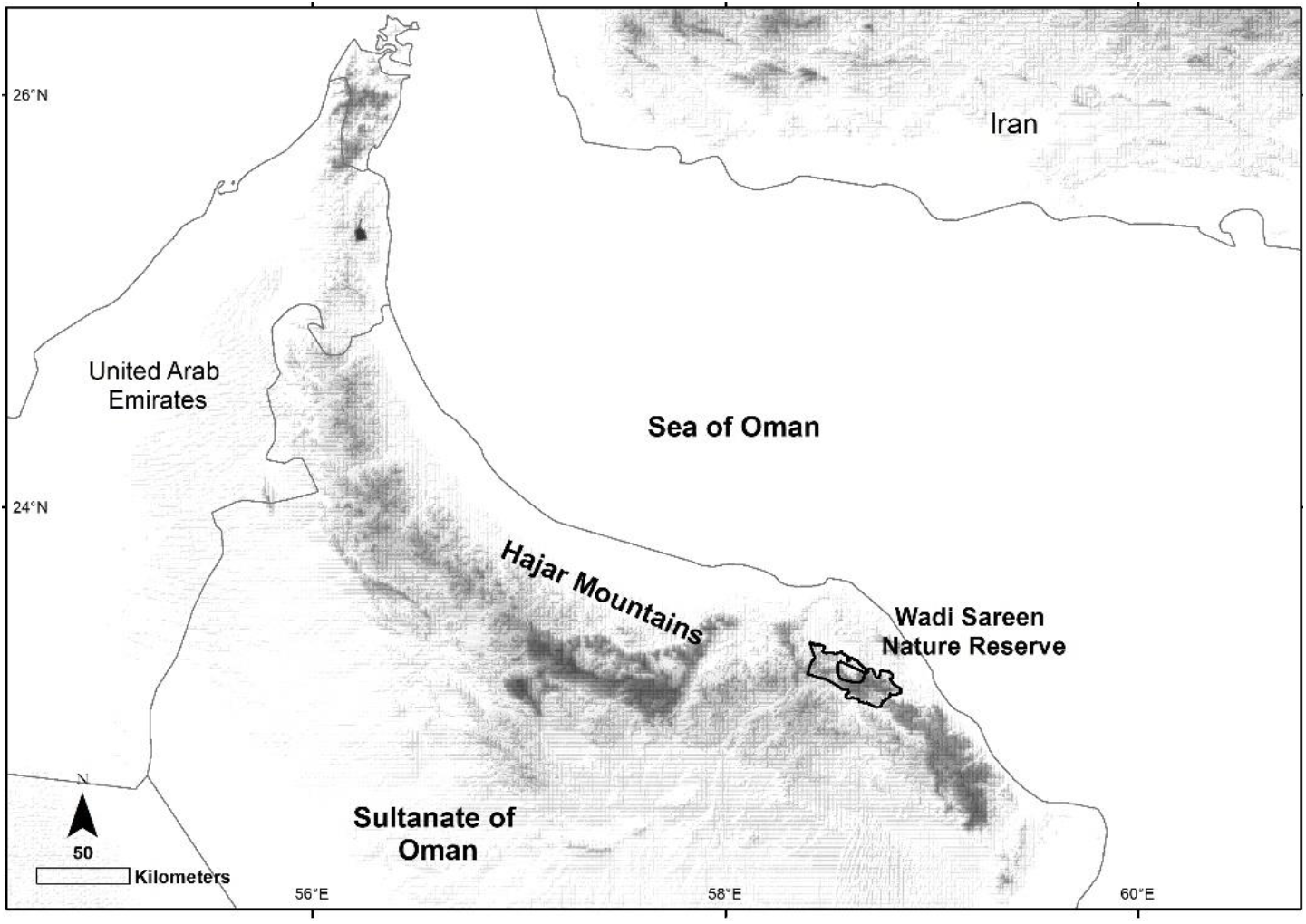
Showing the location of the study area where GPS collaring and tracking of Arabian tahr took place, inside Wadi Sareen Nature Reserve in the Sultanate of Oman.

Considering the Arabian tahr is at risk of extinction, actions need to be taken to secure the species survival. Fundamental to conserving the Arabian tahr is understanding the habitat selection patterns of the species, so that critical habitats can be mapped, protected and managed. Habitat selection studies identify habitats that influence species survival, reproduction and ultimately the species fitness in various habitats (Boyce and Macdonald, 1999). Knowledge and maps of preferred habitats can also guide protected area management and assist in targeting important unprotected habitats at the landscape scale (Morris et al., 2016). Habitat selection can thus be directly applied to facilitating better conservation practices and is particularly important for elusive and threatened species. To improve knowledge of the habitat preferences of Arabian tahr we employed GPS collaring and resource selection functions to understand the species habitat selection characteristics. By comparing model coefficients for habitat selection, we evaluated differences in habitat selection between sexes and seasons and measured the effect of scale on selection patterns.

It has long been understood that habitat selection patterns change depending on the scale at which the analyses are conducted (Boyce et al., 2003; Johnson et al., 2004; Johnson, 1980). Habitats found to be beneficial at one scale may be detrimental at another scale, thus the scale at which we observe animal responses may influence our management decisions (Ciarnello et al., 2007; Hobbs 2003). Investigating resource selection across multiple scales is often used to identify factors that limit species distributions across scales of space and time (DeCesare et al. 2012) and reveals the full repertoire of a species habitat selection behavior. As the distribution and habitat relations of Arabian tahr have been examined at the landscape scale (Ross et al. 2017), we focused on small (foraging) scale and medium (home-range) scales of selection. We aimed to provide managers with habitat selection data at all scales potentially useful for conservation land management.

Gender was also expected to play a role in habitat selection, as sexual segregation is predicted to occur in species when adult males are more than 20% larger than females, as larger size is associated with changes in physiology (Illius and Gordon, 1987). Arabian tahr males are approximately twice the body mass of females (40-45 kg vs. 18-22 kg; Ross, unpublished data), and we hypothesized that this level of dimorphism would result in sexual differences in habitat utilization. In other dimorphic ungulates segregation has been suggested to occur for numerous reasons, including the costs of competition between the sexes (Clutton-Brock and Harvey, 1983), their differing nutritional needs (Demment and Van Soest, 1985), their sensitivity to weather (Conradt et al., 2000), and differences in predation risk, particularly in the presence of young (Main and Coblentz, 1990). Most often a combination of the above reasons influence the fitness of males and females differently, resulting in habitat segregation. At times the differences in habitat selection between sexes of dimorphic ungulates can be large, and gender specific habitat selection becomes relevant to conservation management. In some species of ungulate it has been suggested that male and female ungulates are treated as separate species for purposes of management (Kie and Bowyer, 1999).

Extremely high summer temperatures are the norm in the Arabian Peninsula, and animals must find ways to thermoregulate during high temperatures, physiologically, behaviorally and through their habitat selection patterns (Fuller et al., 2014). As average winter temperatures in Oman are substantially cooler than the summer, seasonal differences in the selection of habitat use for thermoregulation were expected among tahr. In addition, as heterogeneous landscapes have a broader suite of thermal niches for organisms (Elmore et al., 2017), our mountainous study area provided good scope for selection of thermal niche. We aimed to understand how thermal refuges are used as an indication of how Arabian tahr thermoregulate, and how they may use habitat to adapt to future climate changes. In addition, by providing data that allows thermal niche to be mapped we aimed to improve management capacity under climate change.

The primary goals of this paper were therefore to (1) identify key habitat variables contributing to Arabian tahr habitat use, (2) alongside previous large-scale research determine at what scale habitat selection occurs, (3) understand how gender and season influence habitat selection and species management, and (4) identify selection of habitats that provide thermal refuges, to understand potential climate change responses and best management actions.

## Materials and Methods

### Study area and GPS collar data

The study area was located in Wadi Sareen Nature Reserve, in Northern Oman (Fig. 1). The area is typified by steep rocky cliffs and boulder strewn slopes, separated by numerous wadis (ephemeral riverbeds). Elevation ranges from 350 to 1850 m. The study area experiences a hot, arid climate with average maximum and minimum temperatures of 40 and 27.8 C in July, and 22.8 and 13.3 in January, the coldest month. Average annual rainfall in the area is 117.2 mm with most rain falling during the winter (PACA, 2018). Vegetation composition of the study area is largely determined by elevation and water availability, with higher diversity and cover associated with higher altitudes and wadi (drainage) habitats (MacClaren and Rawahi, 2015). Trees typical of the area include *Acacia tortillis, Rhus aucheri, Moringa peregrina*, and *Ziziphus hajarensis*. Common plant species include *Euphorbia larica, Lavandula subnuda, Ochradenus aucheri, Euryops arabicus* among many others.

We captured Arabian tahr at 3 locations on the cliffs of Wadi Sareen between 2012 and 2017, using a custom-made leg snare trap fitted with a trap alarm (Ross and Jahdhami, 2015). All capture procedures were reviewed and permitted by the Ministry of Climate Affairs and the Environment, Muscat. Following capture, tahr were blindfolded, handled without anaesthesia, and fitted with global positioning system (GPS/GSM) tracking collars (GPS-Plus model, Vectronic Aerospace, Berlin, Germany). Position acquisition interval varied from one to six hours and focused on crepuscular foraging activity peaks. The collars collected data for one year before a timed drop-off was activated, the dropped collar was searched for using a VHF beacon, retrieved and downloaded.

### Habitat Selection

We quantified habitat selection using resource selection functions (RSF), where observed GPS points of individual tahr were compared to random available locations (Manly et al. 2002). We modelled the response (used resources) to spatial heterogeneity of available resources at two scales representing 3^rd^ order selection of habitat patches within the home-range or ‘home-range scale selection’, and 4^th^ order selection in buffers surrounding each collected point, highlighting resource selection within habitat patches or ‘foraging scale selection’ (Johnson, 1980).

Due to differences in male and female movement patterns, we calculated available locations for home-range and foraging selection functions separately for each sex. To calculate available points at the home-range scale we measured the maximum distance between all used locations for each individual tahr and defined the buffer radius as the mean maximum distance for males and females (Boyce, 2006). Used points were buffered by 4204 m for females and 5866 m for males. The outer boundary line of all buffered points was used as the area from which available points were sampled randomly at a 1:1 ratio with used points.

We used a case-control design to calculate foraging-scale selection (Compton et al., 2002). Available locations were defined within buffers surrounding each used location and equal to the mean distance travelled between consecutive used locations, calculated separately for males (2572m) and females (1340m). A total of 10 random available locations were generated within the availability buffer around each used location and matched to the used location using a multi-level model design. In this way we constrained available locations to match the scale at which tahr were selecting habitat at each movement step (Johnson *et al*., 2002; Boyce, 2006).

### GIS data and predictor variables

We selected predictor variables using a combination of those shown to influence habitat use in other mountain ungulates, and to reflect the Arabian tahr’s need for food, shelter, predator avoidance and temperature regulation (Lima and Dill, 1990; Mysterud and Ostbye, 1999; Marchand et al., 2014). All variables were measured using a base map including a digital elevation model and satellite imagery within ARCGIS 10.3 software (Environmental Systems Research Institute, Redlands, California). Variables were measured at a 5 m spatial resolution. Variables incorporated into models included ‘elevation’, terrain ‘ruggedness’, which is a measure of topographical heterogeneity and predator cover (Jenness, 2013), distance to escape terrain, ‘dist. to slope’, measured as distance to slopes greater than 35°, dist. to wadi, measured as the distance to a steep sided ephemeral riverbed (wadis) that contain water only under heavy rainfall and mapped using ArcGIS hydrology tools, ‘heat load’, indicating the influence of sunlight on surface temperature, and derived from the slope, aspect and latitude of each site (McCune and Keon, 2002), and the normalised difference vegetation index ‘NDVI’, as an indication of primary production and vegetation availability, and derived from satellite imagery collected in summer and winter seasons of 2014 to 2015. Ground truthing demonstrated that low NDVI signatures were associated with areas of rocky cover and ledges, whereas high NDVI signatures were associated with tree and shrub cover (Maclaren and Rawahi 2015). Variable values were obtained for all used and available points.

### Model construction

Due to large seasonal differences in ambient temperature, positional data were split into summer (April 16^th^ to October 15^th^) when average ambient temperatures were 35 °C and winter (October 16^th^to April 15^th^) when average ambient temperatures were 25 °C (PACA, 2018). Of 26 GPS-collared Arabian tahr, fix data from 24 individuals (10 females, 14 males) were used to derive summer RSF models and 25 individuals (10 females, 15 males) were used to construct winter RSF models.

Models were estimated in MLwiN 2.33, a statistical package for multilevel modelling (Rasbash et al. 2014). To calculate RSFs we used generalized linear mixed-effects models. RSFs included a random intercept to control for the non-independence of individual’s locations and the variable number of observations per individual (Gillies et al. 2006). Habitat variables were used as fixed effects characterising selection. Only variables with a Pearson’s correlation of less than 0.4 were included in the same model. We estimated coefficients of selection (β*i*) to characterize the relative probability of use w(x) using the exponential approximation of the logistic regression model as:

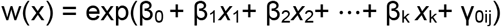

where β_0_ is the intercept for the population and was included because we were using random intercepts (Gillies et al., 2006), β_k_ was the selection coefficient for the k^th^ habitat characteristic and γ_0ij_ was the random intercept for the j ^th^ individual.

Models used a logit link function and Markov Chain Monte Carlo (MCMC) estimation with 100 000 iterations in a single chain. Maximum likelihood parameter estimates starting values were generated using iterative generalized least squares, and the first 5 000 iterations were used as a burn-in for parameter convergence (Spiegelhalter et al. 2002). Four *a priori* candidate models describing tahr habitat selection were run for each season and scale of analyses. *A priori* models included the ‘Risk model’ representing the use of antipredator habitat to avoid predation (ruggedness and dist. to slope), a ‘Climate model’, indicating the use of habitat for behavioural thermoregulation (heat load and elevation), a ‘Vegetation model’, indicting selection in relation to vegetation biomass density (NDVI and dist. to wadi), and a ‘Holistic model’ which was constructed using all variables together except for ruggedness, which was correlated with heat load. We assessed tahr RSF model fit using Diagnostic Information Criterion (DIC; Spiegelhalter et al. 2002), where the model with the lowest DIC score was considered the most parsimonious and models with DIC values differing by > 5 were considered different (Spiegelhalter et al. 2002; Thomas et al. 2006).

We inferred selection or avoidance of resource variables when 95% confidence intervals of fixed effect beta coefficients did not overlap 0. We investigated potential sex-specific patterns in resource selection by including a dummy-coded ‘male’ variable (0 = female, 1 = male) and fitting interactions between gender and each resource variable. We used the Customised Predictions Function in MLwin to explore how the value of variables affected the probability of selection and graphed the selection of individual resource variables (keeping other co-variates constant) of males and females together (Rasbash et al. 2016).

### Model Validation

We used *k*-fold cross validation to test how well habitat selection models predicted selection patterns of the tested population (Boyce et al. 2002). Five folds of cross validation were run, each fold used all data except for 2 withheld individuals. Different individuals were excluded for each fold and used later to test model predictions within ArcGIS (Koper and Manseau, 2012). RSFs for each fold were projected into the ArcGIS database using the models created from the training data sets. The RSF score of each map pixel was reclassified into 10 approximately equal area bins characterizing low to high relative probability of use. A Spearman-rank correlation (Rs) between the area-adjusted frequency of cross-validation points within individual bins and the bin rank was calculated for each cross-validated model. The mean correlation (Rs) across all five folds was used to assess model fit. A model with good predictive performance would be expected to be one with a strong positive correlation, as more use locations (area-adjusted) would continually be falling within higher RSF bins (Boyce et al. 2002). Predictive maps of Arabian tahr resource selection were created using the best cross validated models and mapped using the same range values used for cross validation (Morris et al. 2016).

## RESULTS

Arabian tahr were captured from June to October from 2012 to 2016. A total of 39 animals were captured using a leg-snare trap (Ross and Jahdhami, 2015). Twenty-six Arabian tahr were fitted with GPS collars, 13 captured tahr were not fitted with collars as they were not fully grown and therefore at greater risk of incurring fitness costs of carrying a GPS collar. Three Arabian tahr died while wearing a GPS collar, the cause of death in one male was unknown, one female was observed with severe respiratory issues before dying, with symptoms suggesting Pneumonia, evidence suggested the final male death was due to predation by an Arabian wolf *Canis lupus arabs*.

The variable distance to village was also tested during modelling but not reported as there was no effect shown on Arabian tahr resource selection. This result was of interest as it suggested there is no avoidance response to villages by Arabian tahr inside the protected area.

### Resource Selection, Sexes combined

At the home-range scale the Holistic model was the most parsimonious for both summer and winter seasons, and the best fitting predictive model according to cross-validation (Table 1). At the home-range scale the relative probability of a tahr using a location was positively associated with elevation (Table 2), with higher selection for elevation in the winter (0.224 ± 0.005) than in the summer (0.127 ± 0.005). Tahr also selected areas closer to wadis in summer (−0.191 ± 0.019) and winter (−0.168 ± 0.018), and selected areas closer to steep (>35°) slopes equally in summer (−1.518 ± 0.042) and winter (−1.437 ± 0.036). There was a strong avoidance of locations with high NDVI at the home-range scale, which was a surrogate for tree and shrub cover. This avoidance response was greater in the summer (−1.962 ± 0.198) than in the winter (−0.666 ± 0.180). Tahr also were highly selective of cooler locations with low heat load, with strong selection for lower heat load in the summer (−3.378 ± 0.065) and the winter (−4.016 ± 0.061) when the ambient temperature was lower.

**TABLE 1.** The results of model selection at two spatial scales for Arabian tahr in summer and winter seasons in Wadi Sareen Nature Reserve, Oman. Results include rank according to deviance information criteria (DIC), ΔDIC and an average Spearman-rank correlation (Rs) between area-adjusted frequency of cross-validation points and bin rank for five folds of data.

|  | Home range scale |  |  |  |  |  | Foraging scale |  |  |  |  |  |
| --- | --- | --- | --- | --- | --- | --- | --- | --- | --- | --- | --- | --- |
|  | Summer |  |  | Winter |  |  | Summer |  |  | Winter |  |  |
| Model | Rank | ΔDIC | Rs | Rank | ΔDIC | Rs | Rank | ΔDIC | Rs | Rank | ΔDIC | Rs |
| Holistic | 1 | 0 | 0.95 | 1 | 0 | 0.98 | 1 | 0 | 0.94 | 1 | 0 | 0.97 |
| Climatic | 2 | 3842 | 0.87 | 2 | 3386 | 0.86 | 3 | 4570 | 0.93 | 3 | 7745 | 0.99 |
| Risk | 3 | 5190 | 0.84 | 3 | 4444 | 0.83 | 2 | 100 | 0.97 | 2 | 1677 | 0.96 |
| Vegetation | 4 | 8635 | 0.42 | 4 | 12534 | 0.59 | 4 | 10927 | 0.69 | 4 | 18984 | 0.45 |

**TABLE 2.** Resource selection coefficients for Arabian tahr, sexes combined at two spatial scales, in summer and winter seasons in Wadi Sareen Nature Reserve, Oman.

| Covariate | Home range scale |  | Foraging scale |  |
| --- | --- | --- | --- | --- |
|  | Summer | Winter | Summer | Winter |
| | $\beta \pm SE$ | $\beta \pm SE$ | $\beta \pm SE$ | $\beta \pm SE$ |
| **Intercept** | -1.325 ± 0.083 | -2.527 ± 0.073 | -2.416 ± 0.045 | -2.542 ± 0.035 |
| **Dist_Slope/100** | -1.518 ± 0.042 | -1.437 ± 0.036 | -7.889 ± 0.169 | -9.947 ± 0.188 |
| **Ruggedness** | 1.415 ± 0.090 | 1.705 ± 0.092 | 0.128 ± 0.013 | 0.187 ± 0.011 |
| **Dist_wadi/100** | -0.191 ± 0.019 | -0.168 ± 0.018 | 0.025 ± 0.011 | 0.014 ± 0.010 |
| **NDVI** | -1.962 ± 0.198 | -0.665 ± 0.178 | 1.295 ± 0.210 | -1.454 ± 0.200 |
| **Heat load** | -3.378 ± 0.065 | -4.017 ± 0.061 | -0.749 ± 0.024 | -0.861 ± 0.020 |
| **Elevation/100** | 0.127 ± 0.005 | 0.224 ± 0.005 | 0.004 ± 0.003 | 0.024 ± 0.003 |
| **Random effects** | 0.108 ± 0.036 | 0.055 ± 0.019 | 0.001 ± 0.001 | 0.003 ± 0.002 |

At the foraging scale, the Holistic model was again most parsimonious, although the best fitting predictive model was the Risk model in summer and Climatic model in winter, the top three models all had excellent predictive capability. Similar to the home-range scale, the probability of habitat use increased with elevation, with higher selection for higher elevations in the winter (0.024 ± 0.003) than in the summer (0.004 ± 0.003). Selection for proximity to steep slopes was also maintained at the foraging scale in both seasons (summer −7.889 ± 0.169; winter −9.947 ± 0.188). Tahr maintained their preference for cooler habitat at small scales, with high selection lower heat load in both summer (−0.749 ± 0.024) and winter (−0.861 ± 0.020). Changes occurred in tahr selection of wadis and NDVI. At the foraging scale wadis were avoided in both summer (0.025 ± 0.011) and winter (0.014 ± 0.010). In winter areas of high NDVI were avoided by tahr (−1.454 ± 0.200), but in summer tahr selected locations with higher NDVI (1.295 ± 0.210).

The four best models describing the probabilistic distribution of resource selection across Wadi Sareen Nature Reserve were mapped at the foraging scale (Figure 3) and home-range scale (supplementary materials S1 – S2). At the scale mapped here there is not a large visual difference between maps, however at smaller scales that are more typically used for protected area management, seasonal differences are observable and provide support decision making.

### Sex specific selection

When using sex as an interaction term we found resource selection varied considerably between the sexes (Fig. 2). Both sexes strongly selected for low heat load habitats at home-range scales. However, at foraging-scales males showed significantly stronger selection than females for low heat load habitats, indicating differences in how each sex thermoregulates. Scale and sex specific selection was also evident in the selection of elevation. While both sexes selected higher elevation habitats, females selected more strongly than males at the home-range scale, and males selected higher elevations more strongly at the foraging scale, particularly in summer.

**Figure 2:**
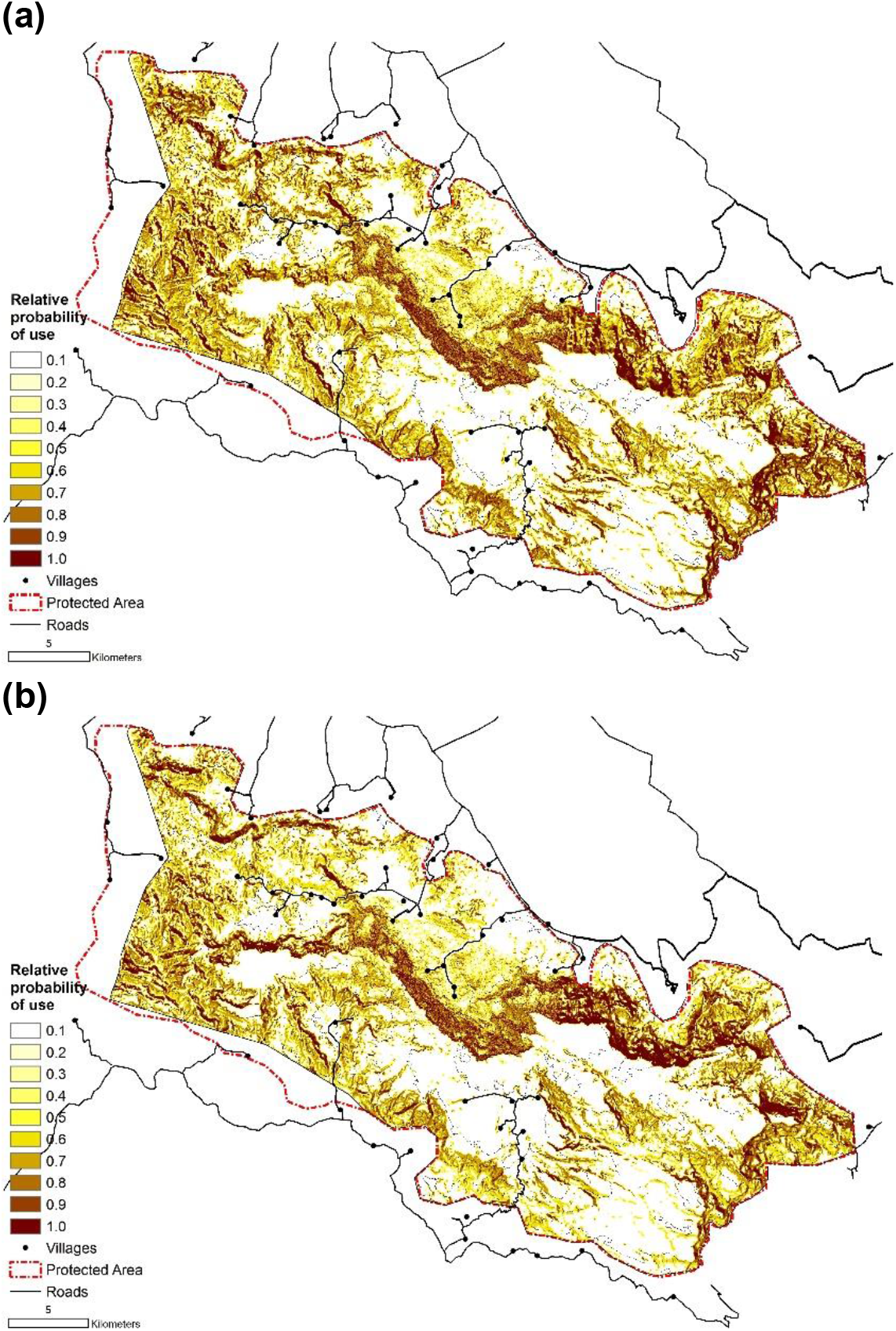
Maps showing the relatively probability of use of Arabian tahr in Wadi Sareen Nature Reserve, Oman, for the most parsimonious foraging scale resource selection models during the (a) summer April 16^th^ to October 15^th^ and (b) winter October 16^th^to April 15^th^.

**Figure 3:**
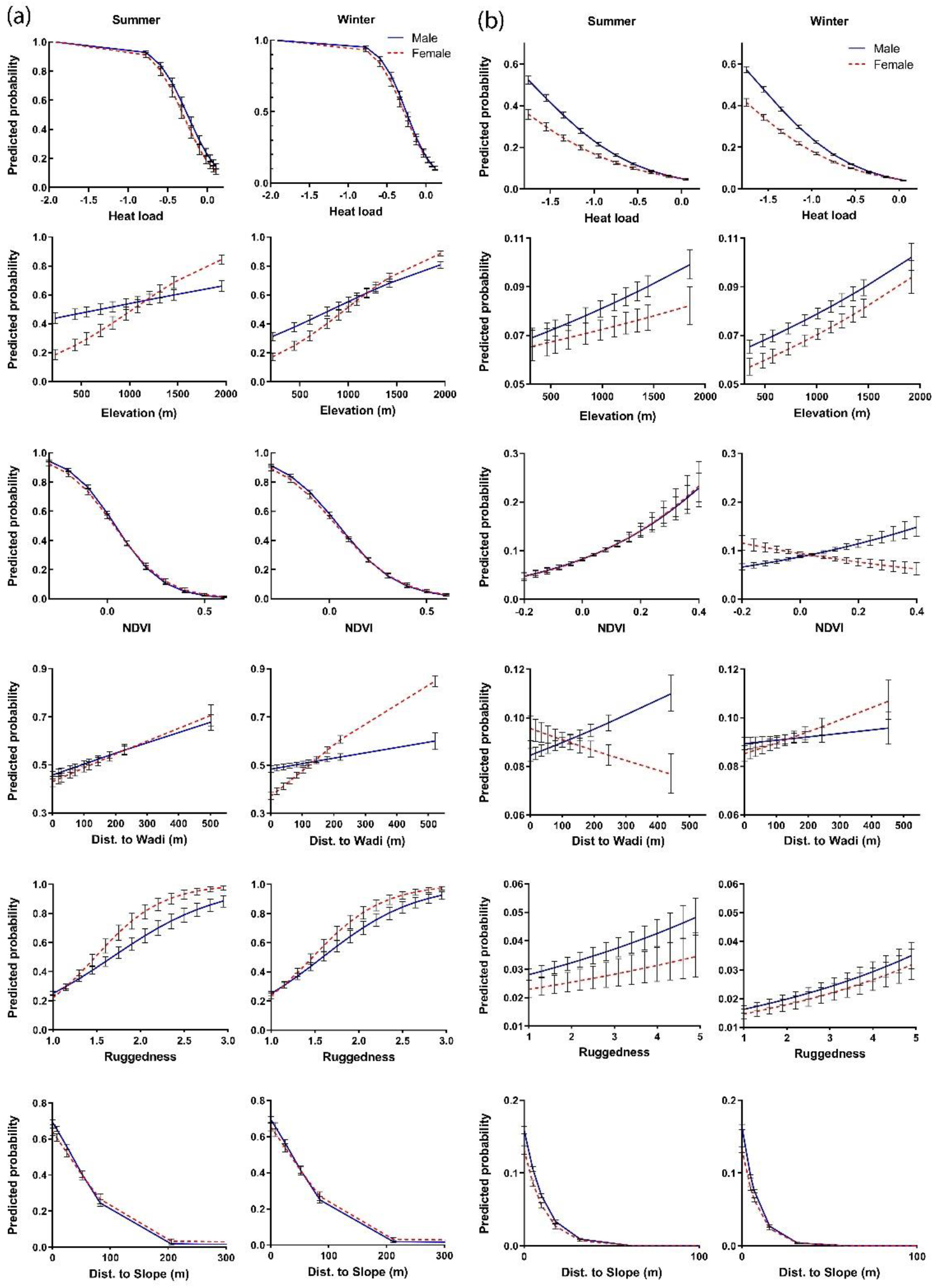
Response of Arabian tahr resource selection probability (predicted probability) to increasing key variables, keeping other model variables constant, at the home-range scale (a) and foraging scale (b), in summer (left) and winter (right). Graphs show the male (solid) and female (dashed) responses, error bars indicate the 95% error of individual variability.

Male and female selection of rugged and steep habitats were similar for both seasons, but females showed higher selection for rugged habitats at home-range scales and males selected rugged and steep habitats more strongly at the foraging scales.

Tahr showed scale and sex specific selection of NDVI. Areas with higher NDVI were avoided at the home-range scale by both sexes, but at foraging scales sexual differences were apparent. In winter males selected habitats with higher NDVI at foraging scales whereas females preferred lower NDVI, less vegetated habitats, suggesting a preference for rocky and sparsely vegetated habitats. In summer both sexes had equally strong selection for habitats with high NDVI, suggesting a focus on habitats with more tree and shrub cover. In both seasons wadi habitats were selected at the home-range scale. At foraging scales both sexes avoided wadis in winter, males maintained their avoidance for wadis the in summer, but females showed selection for habitats closer to wadis in the summer season.

## Discussion

We examined the habitat selection patterns of the Arabian tahr and how sex, scale and season affect habitat preferences. Our study demonstrated that sex, season, and scale influence resource use and are important considerations in the conservation management of tahr. The new understanding of resource use is important to preserving the integrity of critical habitat for Arabian tahr and managing availability of habitats that contribute to the species resilience to climate change.

As conservation management is practiced at multiple scales, the scale at which an animal selects resources can guide the management and protection of species and their habitats. For example, feeding resources of ungulates are generally selected at very small scales, whereas selection of habitats used to evade predators are usually selected at larger scales (Creel et al., 2005; Johnson, 1980). Large-scale selection of anti-predator habitat allows ungulates to forage for food safely and efficiently within the selected home-range area. Scale-specific selection patterns were also evident among tracked Arabian tahr. Variables related to food density, such as NDVI, were highly selected at the smallest foraging scale of analyses but were avoided at larger (home-range) scales of selection. Similar findings were also found beyond the home range area, for example Ross et al. (2017) found that NDVI did not predict tahr occupancy at the landscape scale. Questions relating to the management of food resources for Arabian tahr are thus best addressed at small scales and are likely to be missed by larger scale studies.

Given the high temperatures experienced in the study area it was not surprising that thermal refuges with lower heat loads were highly selected at both foraging and home-range scales. Although large scale research found no association between heat load and tahr presence (Ross et al. 2017), heat load selection at the home-range scale suggested tahr consider temperature when selecting home-range areas for foraging, and again while they are in the process of foraging. Although past literature suggests that thermal refuges are commonly patchy and of relatively small size (Alston et al., 2020), our research shows that selection of habitat providing thermal refuge may also be more premeditated and occur at larger spatial scales.

In contrast to food and temperature requirements, we found rugged habitats, higher elevations and steep slopes were consistently selected across all scales tested here and in previous large-scale studies (Ross et al., 2017). The selection of rugged, elevated, and steep habitats across scales suggests these habitat characteristics are core requirements of Arabian tahr. As caracal *Caracal caracal*, Arabian wolf *Canis lupus arabs* and occasionally poachers were found in the study area, rugged and steep habitats are most likely selected to provide important antipredator escape terrain (Rachlow and Bowyer, 1998; Bangs et al., 2005). However, shade and productive microclimates for herbaceous plant growth were also found in rugged terrain and likely influenced selection. The consistent selection of higher elevations, across scales, suggests preferences for lower temperatures, greater incidence of fog, and high-altitude plant associations (Maclaren and Rawahi, 2015).

As well as scale we were interested in how sex affected selection, and whether gender is an important factor in tahr management. Some form of sexual segregation was expected to occur as male tahr are 50% larger than females, which are likely to result in differences in physiology (Illius and Gordon, 1987). Due to larger body size and longer food retention times male tahr should be able to subsist on a lower quality, more fibrous diet than the smaller female tahr (Demment and Van Soest, 1985; Illius and Gordon, 1992). Larger males should also have a relatively lower perturbation of core body temperature when exposed to high temperatures (Jessen, 2001; Fuller et al. 2016), which could lead to male tahr having less need for low heat load, thermal refuges.

Following predictions, sexual segregation was indicated in the selection of habitats associated with food. Though avoided at the home-range scale, NDVI selection occurred at the foraging scale, with males selecting habitats with higher NDVI year-round, and females selecting high NDVI habitats only during the summer period. Given males ability to subsist on a more fibrous diet than females (Illius and Gordon, 1992), their year-round small-scale selection of tree and shrub cover may have optimized their biomass and nutritional intake. Females avoidance of tree and shrub cover during winter most likely related to a preference for rocky habitats and ledges which have low tree and shrub cover but good cover of grasses and herbaceous plants during the winter season. In the hot summer female selection of shrub and tree cover may relate to a higher nutrient content than grass and herb cover (Gordon and Illius, 1989; Gordon and Prins, 2008), grasses and herbs were also very scarce in the summer. Both sexes may also take advantage of the numerous trees and shrubs that bear fruit and seeds in the study area in the summer (e.g. Ghazanfar, 2018). Wadis were generally avoided by both sexes, but females selected wadis in summer, which could relate to greater nutrition found in the drainage channels, which tend to remain greener over the summer, but may also be used by females to provide cover for young, which are born in April and May. Wadis in the area are full of large boulders and crevices and young tahr have been observed hiding in the boulder strewn wadis (S. Ross, unpublished data). It is possible that wadis acted as a safe creche habitat for Arabian tahr in the study area.

Both sexes strongly selected habitats offering thermal refuge in summer and winter, however the intensity and scale of selection for thermal refuges differed between the sexes, suggesting subtle differences in how the sexes thermoregulated. Microclimate selection is one of the primary means employed by many species to buffer changes in ambient temperatures (Fuller et al., 2014). As temperatures often exceeded 45°C in the study area and remained hot year-round, it was no surprise that both sexes had strong selection for cooler habitats and higher elevations. However, males more intensively selected shade habitats and higher elevations at the foraging scale, whereas females showed higher selection for elevation than males at the home-range scale. The separation between males and females occurred in both seasons but was greater during the summer. The patterns suggest that by more strongly selecting higher elevations at larger scales females prioritize cooler temperatures at a more primary level and therefore have a lower requirement for shade at the foraging scale. Whereas males deal with their weaker selection for elevation at the home-range scale by more strongly selecting shady and more elevated habitats in their daily foraging movements. While males selection of higher elevation at foraging scales are unlikely to result in substantially lower temperatures, selecting local elevated peaks and ridges while foraging most likely provide conductive cooling through wind exposure. The need for shady habitats did not appear to increase in the summer when temperature was considerably higher. However, male and female tahr selected high NDVI habitats during the hot season. Their preference for tree and shrub cover fulfills a need for food and nutrition, but as trees also provide shade, high NDVI habitats were also thermal refuges, particularly during the summer.

## Conclusions

Our study demonstrated that Arabian tahr resource needs differed depending on the scale examined. The scale-specific responses found by our study support previous research suggesting that decisions based on only one scale of analysis are limited in their scope (Kotliar and Wiens, 1990; Guisan and Thuiller, 2005; Ciarnello et al. 2007). At the scale of protected area management, thermal refuges and vegetation were particularly important. As gender and season influenced the response to these variables, managing to support seasonal and gender specific requirements should be considered. Within protected areas actions to cover small scale needs are best managed by providing sufficient undisturbed habitat that allows connections between summer and winter habitat patches. Larger landscape connectivity is also required for potential adaptive range shifts due to climate change. Supporting this need for connectivity, our study indicated that unlike villages outside of Wadi Sareen, villages inside the protected area were not avoided by Arabian tahr. Maintaining this low human impact, by preserving good community relationships, is important for the future ecological integrity of the area.

Certain habitats were found to be equally important at both small and large scales and were less affected by gender or season. Following previous authors (Ciarnello et al. 2007), we believe that habitats selected across scales represent resources of fundamental importance to Arabian tahr. Rugged, steep and high elevation habitats are among the habitats that should be prioritized at all management scales. While rugged and steep habitats are relatively safe from impacts due to the difficulty of accessing them, elevated habitat is a limited resource that is likely to be important in the future to enable climate change adaptation.

Thermal refuges were identified as a critical habitat of Arabian tahr, allowing the species to thermoregulate despite typically high ambient temperatures. The importance of behavioural thermoregulation to tahr is of increasing relevance considering predicted climate change in the region (Pal and Eltahir, 2016). Authors have stressed our need for a spatio-temporal understanding of thermal cover for conservation planning and management actions (Elmore et al., 2017). Our study showed that thermally favorable habitats are selected both at home-range and the foraging scale, suggesting managers should focus on these scales when considering thermal refuges. Our mapping of heat load and elevation allowed us to understand selection of thermal refuge, and similarly can be mapped and visualized by protected area managers to assist in decision making. Climate change is likely to result in preferred foraging habitats shifting to higher elevations. The high altitude of Wadi Sareen Nature Reserve gives scope for movement up slope, with approximately 500-700 m of upwards migration possible, which could help buffer climate impacts. Although habitats may not be of equal quality at higher altitudes, and due to the conical shape of mountains habitat availability generally decreases with increasing elevation (e.g. Lovari et al., 2020). Thus, an overall decline in both habitat availability and suitability is possible in the long-term at all scales. This is of more immediate concern to tahr populations living at lower elevations such as the important Jabal Qahwan population. With little scope for upward migration and due to population isolation (Ross et al., 2020), Arabian tahr in Jabal Qahwan are at particular risk from climate change.

Given our uncertainty of how to conserve wildlife under future climate change, understanding the availability and distribution of wildlife habitat is an important baseline from where we can plan, connect and preserve the resources necessary for wildlife adaptation and conservation.

## Acknowledgements

This study was funded and facilitated by the Diwan of Royal Court, Oman, National Field Research Centre for Environmental Conservation, the Office for Conservation of the Environment, and by the Oman Earthwatch Program. We thank Dr Saif Al Shaksi, Yasser Al Salami, and Dr Mohammed Al Balushi for providing permissions and support for the project, and Dr Roderic Dutton and Nigel Winser for their direction and oversight. We thank the rangers of Wadi Sareen Nature Reserve for their support during animal trapping and fieldwork.

## Author contributions

SR conceived the research, undertook animal trapping and collaring, data analyses and co-wrote the article. MHAJ and HAR assisted with animal trapping and co-wrote the article.

## Notes

### Competing Interest Statement

The authors have declared no competing interest.

