## Supplemental figure S1 and S2 for "Resource selection by the Endangered Arabian tahr: identifying critical habitats for conservation and climate change adaptation"

(a)

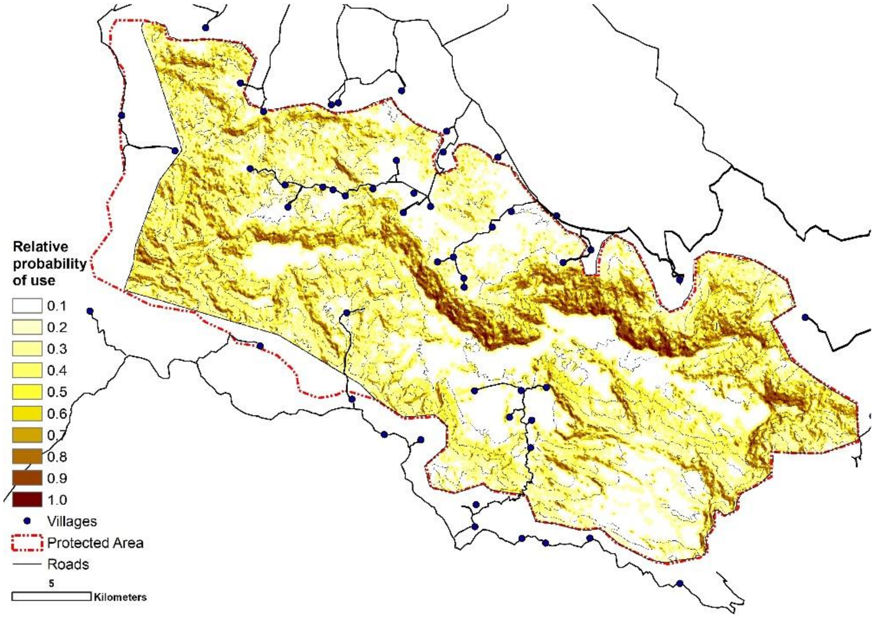


(b)

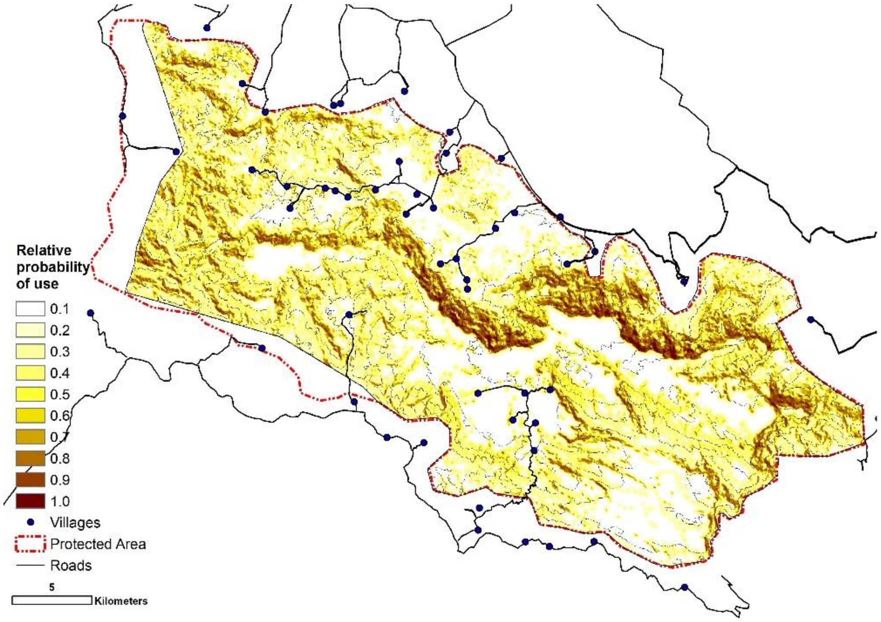


Figure S1 and S2: Maps showing the relatively probability of use of Arabian tahr in Wadi Sareen Nature Reserve, Oman, for the most parsimonious home-range scale resource selection models during the (a) summer April 16th to October 15th and (b) winter October 16thto April 15th.
